# AI-like parallelogram geometry of word relations in the human neocortex

**DOI:** 10.64898/2026.09.07.749996

**Authors:** Ruining Wang, Yujie Tian, Keiko Ohmae, Shogo Ohmae

## Abstract

The same semantic relation (e.g., queen−king and woman−man) is encoded as a shared vector direction in language AI, forming parallelogram geometry, whereas its presence in the human brain remains unknown. Our fMRI analysis revealed similar geometry in the neocortex, particularly in higher-order language regions. Our findings link the brain and AI through a common geometry, advancing understanding of how abstract relational knowledge is organized and used in human language processing.

---

Recent advances in language AI have created a new opportunity to understand human language processing by directly comparing internal representations in artificial systems and the human brain, thereby revealing intriguing brain-AI parallels and convergence (as recently reviewed^1^). Such parallels have been identified across multiple levels of linguistic representation, from words and phrases to syntactic structure and sentences^2-7^. At the level of word meaning, a central organizing principle shared by the cerebral neocortex and language AI is categorical organization, in which semantically similar words cluster together in representational space, as observed in fMRI mapping^8^ and single-neuron recordings^9^.

Beyond categorical organization, language AI exhibits another geometric property of word representations that has not been directly examined in the human brain. In artificial word representations, difference vectors encoding the same semantic relation, such as “queen − king” and “woman − man,” tend to be parallel, forming a parallelogram among the four words (**Fig. 1a**). This parallelogram represents the semantic relation itself as a common vector direction, enabling analogy through linear vector operations such as *queen* ≈ *king* − *man* + *woman* and contributing to the efficient organization of tens of thousands of words in a relatively low-dimensional space^10-15^. Although this structure has been widely observed from early word embeddings to recent Transformer-based language models, whether a similar relation-vector geometry exists in word representations in the human brain remains unknown. Notably, in language AI, this geometry is more prominent in intermediate layers than in input layers^10, 13, 16^. We therefore examined whether word representations in the human brain exhibit similar parallelogram geometry, and whether its strength increases along the cortical hierarchy of language processing.

**Figure 1.**
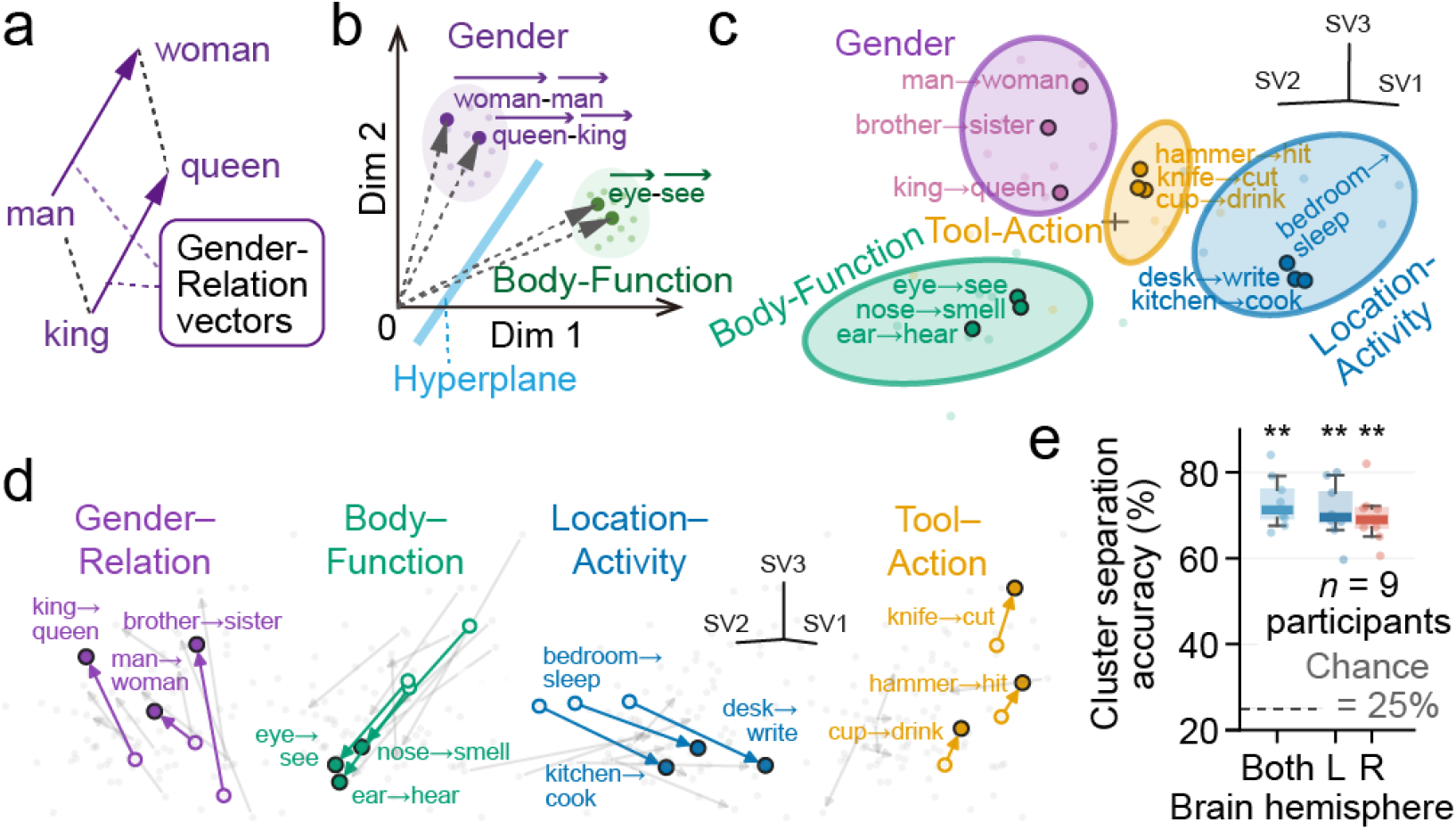
Word-relation parallelogram geometry in the human neocortex. **a**, Schematic illustration of relation-vector geometry. Word representations form a parallelogram as difference vectors representing the same semantic relation point in similar directions. **b**, Schematic of clusters of recentered relation vectors. Cluster separation was evaluated by a classifier hyperplane. **c**, Relation-vector clusters. Recentered relation vectors from a representative test set (3 pairs) were displayed in the three linear SVM axes (SV1–SV3). **d**, Relation-vector parallelogram geometry. Relation vectors for the same test set were displayed in the same 3D space without recentering. **e**, Cross-validated cluster separation accuracy for the whole brain and the left and right hemispheres (across-participant median, interquartile range, and 5th-95th percentile; one-sided Wilcoxon signed-rank test, W = 45, 45, and 45, *P* = 0.002, 0.002, and 0.002). **, 0.001≤ *P* < 0.01.

To address the first question, we extracted word representations from fMRI data acquired while participants listened to natural narratives and analyzed relation vectors for four semantic relation groups: gender, body–function, tool–action, and location–activity. Recentered relation vectors (**Fig. 1b**) formed clear clusters according to semantic relation (**Fig. 1c**), with cluster separation accuracy reaching 72.9% (median; *P* = 0.002; **Fig. 1e**). In the word representations before recentering relation vectors, sets of four words sharing the same semantic relation formed near-parallelogram configurations (**Fig. 1d**). Therefore, word representations in the human brain exhibit a parallelogram geometry in which semantic relations are represented as shared directions.

Regarding the second question, we observed a clear hierarchical gradient in the strength of this geometry. Cluster separation accuracy for relation groups was higher in higher-order language and association areas than in sensory and motor regions (*P* < 0.001; **Fig. 2a**). A voxel-wise analysis of contributions to cluster separation showed that voxels with high contributions were concentrated in higher-order language and association areas (**Fig. 2b,c**). Thus, as in language AI, the geometry of word relations was more prominent at higher levels of the hierarchy of language and semantic processing.

**Figure 2.**
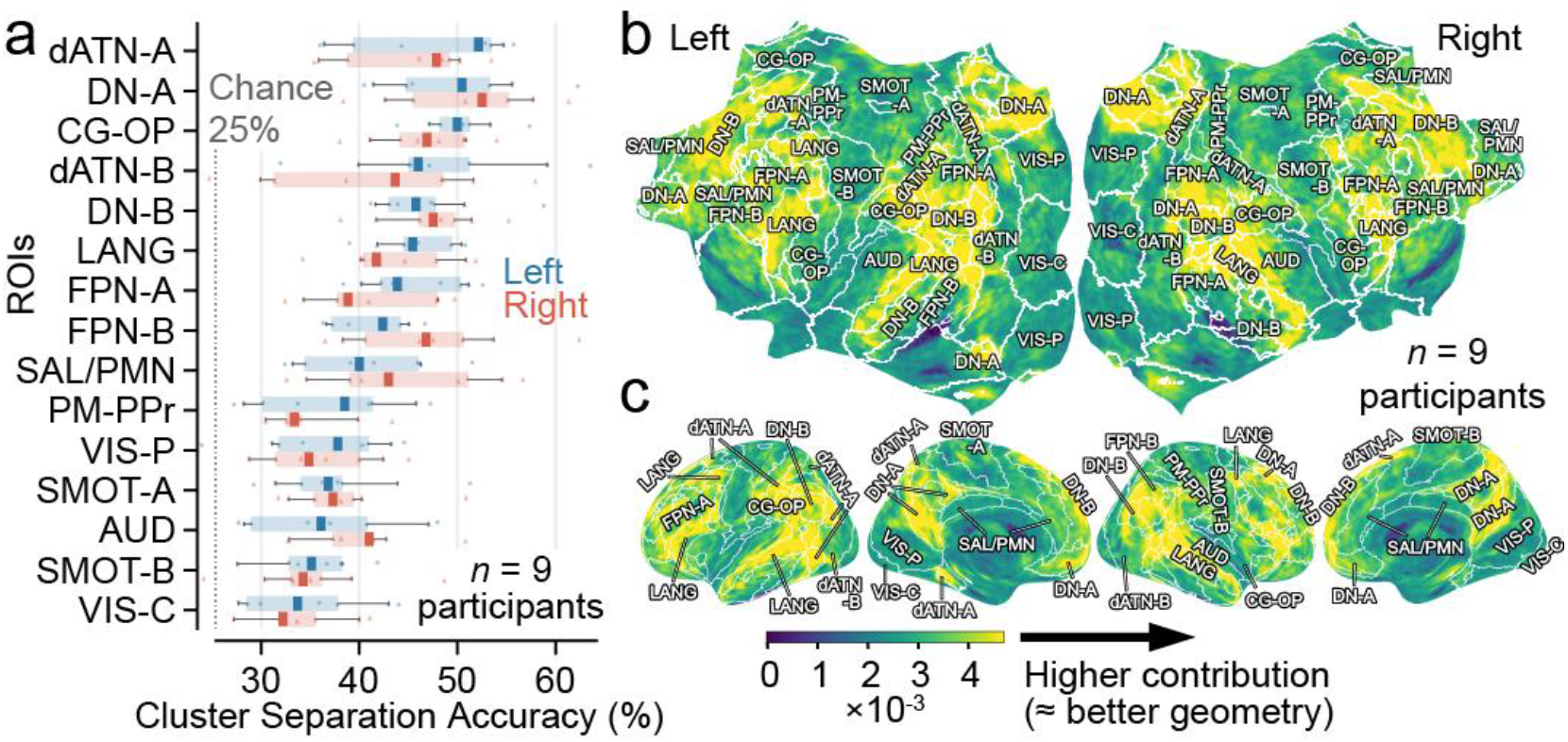
Neocortical distribution of relation-vector geometry strength. **a**, Cluster separation accuracy of relation vectors across 15 neocortical functional network ROIs in the left and right hemispheres (median, interquartile range, and 5th-95th percentile). The accuracies were significantly different across ROIs (Friedman test, χ^2^(14) = 75.99, *P* < 0.001). **b**,**c**, Voxel contribution to the cluster separation displayed on flattened (**b**) and partially flattened (**c**) brain maps. Contributions were derived from linear SVM classifier weights and averaged across 9 participants. AUD, Auditory; CG-OP, Cingulo-Opercular; dATN-A/B, Dorsal Attention-A/B; DN-A/B, Default Network-A/B; FPN-A/B, Frontoparietal Network-A/B; LANG, Language; PM-PPr, Premotor-Posterior Parietal Rostral; SAL/PMN, Salience/Parietal Memory Network; SMOT-A/B, Somatomotor-A/B; VIS-C, Visual Central; VIS-P, Visual Peripheral.

We next quantified this geometry using a measure that does not depend on the number of relation groups examined. Cosine similarity between relation vectors within the same semantic relation revealed significant parallelism in the whole neocortex (cosine similarity = 0.26, mean across 9 participants, two-sided one-sample t-test, *P* < 0.001; **Fig. 3a**), and its neocortical distribution was consistent with that in **Fig. 2** (**Fig. 3b**; FDR-adjusted *P* < 0.05 in most of ROIs, Extended Data Fig. 2b).

**Figure 3.**
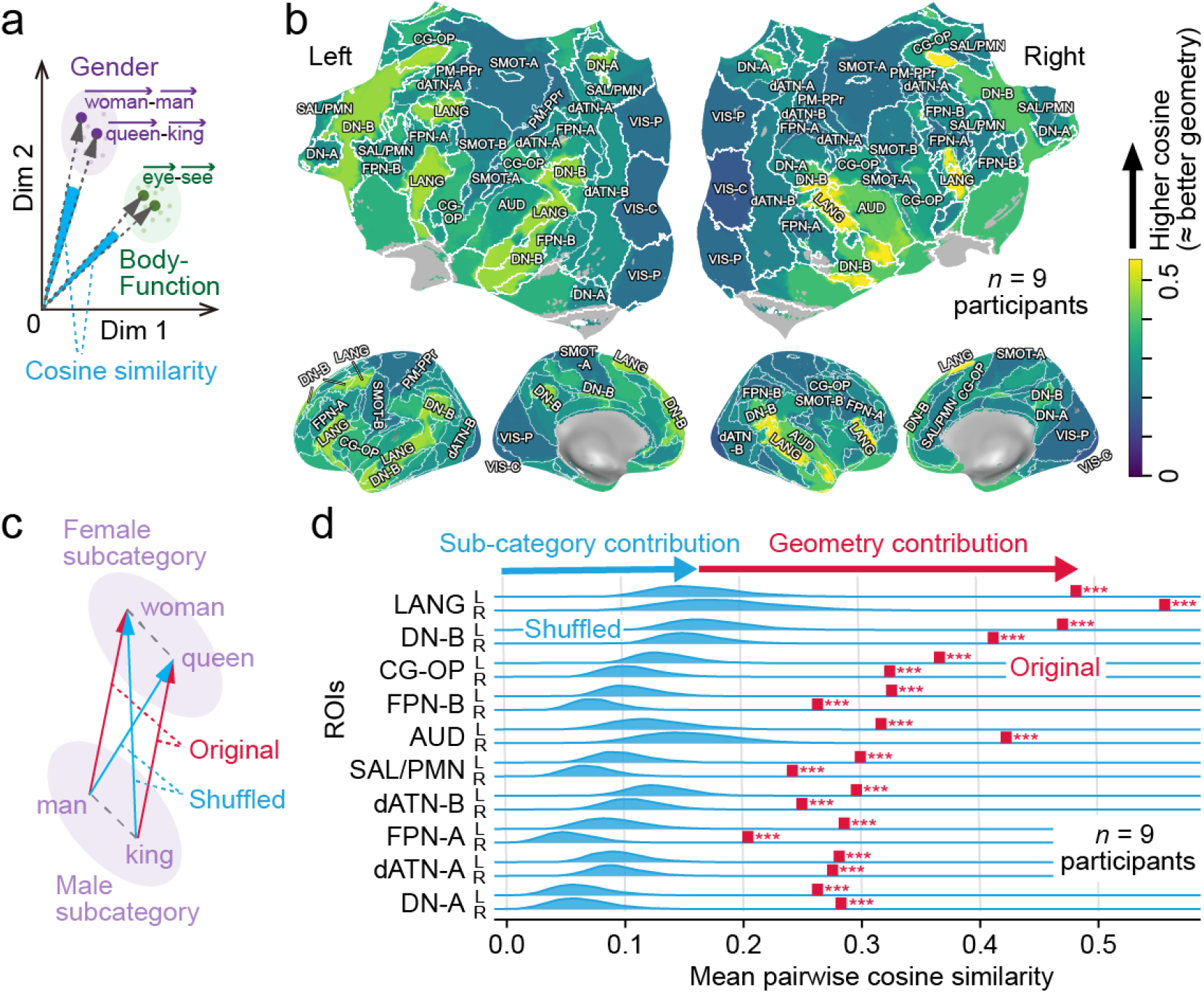
Direct measurement of relation-vector parallelism and subcategory-preserving control. **a**, Schematic illustration of relation-vector parallelism measured directly by cosine similarity between relation vectors belonging to the same relation group. **b**, ROI-wise relation-vector cosine similarity of the gender group. **c**, Schematic illustration of the potential contribution of semantic subcategory separation to apparent parallelism and subcategory-preserving shuffle control (blue). **d**, Cosine similarity of the shuffled and original relation vectors of the gender group (two-sided permutation test, B = 10,000, with Benjamini–Hochberg FDR correction). ***, FDR-adjusted *P* < 0.001.

Finally, we tested a critical alternative explanation: that the apparent parallelism simply reflected subcategory separation. For example, male and female words forming distinct subclusters could produce apparently parallel relation vectors (**Fig. 3c**). However, shuffling the word pairings while preserving the subcategories substantially reduced parallelism (FDR-adjusted *P* < 0.001 in all ROIs; **Fig. 3d**). Thus, the observed parallelism reflected the geometry of semantic relations itself, not merely the arrangement of subcategories in representational space.

This study demonstrated that word representations in the human brain exhibit a parallelogram geometry similar to that in AI. Previous studies have shown neocortical representations corresponding to semantic relations and decoding of target words through vector addition and subtraction^17, 18^, but did not directly establish the parallelogram geometry itself. A recent preprint independently reported aligned semantic-relation directions and parallelogram structures in single-neuron populations in three human brain regions^19^. The present study independently established this word-relation geometry, and further ruled out an alternative explanation based on subcategory separation alone (**Fig. 3c,d**). Our study also mapped the neocortex-wide distribution of this geometry, revealing that it is most pronounced in higher-order language and association areas, in line with recent reports that increasingly abstract and context-dependent language representations emerge in these areas^3, 4, 20, 21^.

Why does this parallelogram geometry emerge in word representations in the brain? This geometry is neither unique to word relations nor confined to a particular system: it appears in nonlinguistic domains in monkey and human brains^22, 23^ and across AI systems with different architectures and learning mechanisms^10, 12, 16^. Thus, relation-vector parallelism may reflect a general representational principle by which relations recurring across different pairs are abstracted into a common direction. Word relations in the human brain may represent one instance of this principle.

What might the brain use this geometry of word relations for? Neuroscience studies have linked parallel geometry to inference and generalization^22, 23^. In AI, operations on semantic relation vectors support analogy, and parallel representations facilitate generalization and efficient learning^14, 24^. Together, these findings suggest that parallel geometry of word relations supports analogy, generalization, and efficient learning in human language processing.

Our findings link the brain and AI through a common geometric structure, opening a new avenue toward understanding how abstract relational knowledge is organized and used in language processing, and how it ultimately supports human intelligence.

## Acknowledgments

This work was supported by Beijing Key Laboratory of Brain Science and Brain-Machine Interface.

## Author Contributions

Conceptualization, S.O.; Methodology, S.O., R.W. and Y.T.; Software, R.W. and Y.T.; Validation, Y.T. and R.W.; Formal analysis, R.W. and Y.T.; Investigation, R.W. and Y.T.; Data curation, Y.T.; Visualization, R.W., S.O., Y.T. and K.O.; Resources, S.O.; Supervision, S.O.; Project administration, S.O.; Funding acquisition, S.O.; Writing – original draft, S.O. and K.O.

## Methods

### fMRI dataset and word representations

We used an fMRI dataset acquired while participants listened to natural stories^8^. Our analyses were conducted on data from nine participants (UTS01-09)^25^. Preprocessing of the fMRI data and estimation of voxel-wise responses to each of the 985 words followed the methods and codes of Huth et al^26^.

In the whole-brain analysis, each word was represented as a high-dimensional word-representation vector consisting of approximately 90,000 voxels. To reduce computational cost and standardize the dimensionality of the analyses, principal component analysis (PCA) was applied to the whole-brain word representations of each participant, reducing them to 46 dimensions. These 46 components explained approximately 70% of the variance in whole-brain word responses (in participant-native space, 58.8–72.1%; in MNI space, 70.6–83.2%). The same 46-dimensional PCA space defined in the whole-brain analysis was also used for the ROI analyses. Specifically, voxel signals within each ROI were projected into the same 46-dimensional space before further analysis. All subsequent analyses were performed independently for each participant.

For across-participant comparison or averaging (**Figs. 2, 3**), word representations were transformed into MNI space before further processing.

### Ethics

The original data collection was approved by the Institutional Review Board of the University of Texas at Austin (protocol no. 2017-07-0030), and written informed consent was obtained from all participants for both participation in the research and public sharing of their data. The present study involved secondary analysis of publicly available, de-identified data and no new participant recruitment or data collection.

### Word pairs and relation vectors

To examine semantic relationships between words, we selected 47 word pairs from the 985 words and classified them into four semantic relation groups. The gender relation (male-female), body– function, tool–action, and location–activity groups contained 12, 11, 12, and 12 pairs, respectively. For each word (w), its 46-dimensional brain-activity vector was denoted as **w**, and the relation vector between a source word (**s**) and a target word (**t**) was defined as **t** – **s** (**Fig. 1a, b)**.

Gender pairs include (12 pairs): man-woman, king-queen, father-mother, son-daughter, uncle-aunt, brother-sister, grandfather-grandmother, gentleman-lady, he-she, his-hers, him-her and male-female.

Body-Function pairs include (11 pairs): eye-see, ear-hear, nose-smell, mouth-speak, foot-walk, leg-run, hand-touch, finger-point, arm-carry, head-think and skin-feel.

Tool-Action pairs include (12 pairs): knife-cut, hammer-hit, gun-shoot, car-drive, boat-sail, net-catch, sword-fight, ladder-climb, radio-listen, television-watch, telephone-call and cup-drink. Location-Activity pairs include (12 pairs): bedroom-sleep, kitchen-cook, library-read, restaurant-eat, office-work, bridge-cross, shop-buy, pool-swim, school-learn, park-play, bath-wash and desk-write.

### Cluster separation analysis of relation vectors

We used linear support vector machines (SVMs) to assess whether relation vectors form clusters according to their semantic relation groups.

Cluster separation was performed using a hierarchical classifier consisting of three linear SVMs. The first classifier separated location–activity and tool–action groups from body–function and gender groups. The second classifier separated location–activity and tool–action groups, and the third classifier separated body–function and gender groups. Each classifier was trained in the 46-dimensional recentered relation-vector space (**Fig. 1b**). The three linear SVMs were trained using only the training pairs. Using this hierarchical classification, the test set was ultimately classified into one of the four semantic relation groups. The chance level for four-class classification was 25%.

To quantify cluster separation accuracy, cross-validation was repeated 400 times. In each repetition, to balance the sizes of the training and test sets, we sampled an equal number of pairs from each group (e.g., eight training and three test pairs; four-fold cross-validation). The hierarchical linear SVM was trained using only each training set, and cluster separation accuracy was calculated for the balanced held-out test set. The accuracies were averaged across 400 repeats, for each participant (**Fig. 1e**). The regularization parameter of the SVM was set to 0.001. For visualization of a representative example, the perpendicular vectors defining the classification hyperplanes of the three SVMs were denoted as SV1, SV2, and SV3 in the figure, and the relation vectors were projected onto these three axes to obtain a three-dimensional representation (**Fig. 1c**). Test pairs were not used to determine these SVM axes.

### Relation-vector geometry across cortical networks

To examine how relation-vector geometry differed along the cortical hierarchy of language processing, the same cluster separation analysis was performed separately for 15 cortical functional network ROIs (**Fig. 2a**).

For each ROI in each hemisphere, the analysis was restricted to voxels within the ROI, and the word representations were projected into the same 46-dimensional PCA space defined in the whole-brain analysis. The resulting relation vectors were then subjected to the same hierarchical linear SVM classification with 400-repeat cross-validation as in the whole-brain analysis.

In **Figure 2a**, the networks were ordered according to their median cluster separation accuracy in the left hemisphere. Differences in accuracy across ROIs were assessed using the Friedman test.

### Voxel contribution to relation-vector cluster separation

To identify cortical regions contributing to the separation of relation-vector clusters, we calculated the voxel-wise contribution to cluster separation using the classifier weights of the linear SVMs (**Fig. 2b, c**).

For each voxel, the coefficient vectors of the three hierarchical SVM classifiers, SV1–SV3 (normal vectors to the separation hyperplanes), were back-projected from the PCA space onto the corresponding dimension of the original voxel space, representing that voxel’s activity. To equally evaluate the contribution of each of the three hierarchical classifiers, the resulting voxel-weight vector was normalized such that the total L2 norm across all voxels for each classifier was one. The results were nearly identical without this normalization (Pearson’s *r* = 0.983). The normalized voxel-weight values were then combined across the three hierarchical classifiers by calculating the square root of the summed squared weights and averaged across 400 repeats to generate the voxel-wise contribution value. Finally, the contribution values were averaged across the nine participants in common MNI space and displayed on a flattened cortical surface (**Fig. 2b**) and a partially flattened cortical surface (**Fig. 2c**).

### Direct measurement of relation-vector parallelism

As a geometric measure independent of classification, we directly evaluated vector parallelism using the cosine similarity between relation vectors belonging to the same semantic relation group (**Fig. 3a,b**). To remove apparent parallelism (high cosine similarity) arising from a component shared across all relation vectors, we subtracted the mean relation vector across all word pairs from each individual relation vector. Cosine similarity was calculated for all distinct combinations of word pairs within each semantic relation group, and the mean was used as an index of parallelism. Higher cosine similarity indicates that relation vectors point in more similar directions. To facilitate comparison with the classification analysis in **Figure 2**, the cosine similarity for each ROI was plotted on a brain map (**Fig. 3b**).

### Subcategory-preserving shuffle analysis

To test whether the observed relation-vector parallelism could arise just from the separation of semantic subcategories of the source and target words, rather than from the semantic relation geometry itself, we performed a subcategory-preserving shuffle control (**Fig. 3c,d**).

For example, in the gender relation, source words belong to the male subcategory and target words to the female subcategory. Even if male and female words simply form separate clusters, the relation vectors connecting them could show apparent parallelism (**Fig. 3c**). We therefore randomly shuffled only the correspondences between words within each subcategory while preserving the source-side and target-side subcategories.

Relation vectors were recalculated from the shuffled pairs, and cosine similarity was calculated in the same manner as for the original pairs. By comparing parallelism between the original and shuffled data within participants, we assessed the component of the relation-vector geometry that could not be explained solely by subcategory separation (**Fig. 3d**).

### Statistical analysis

Whole-brain cluster separation accuracy was compared with the chance level of 25% using a one-sided Wilcoxon signed-rank test. Differences in cluster separation accuracy across network ROIs were assessed using the Friedman test.

Cosine similarity of the whole brain was tested against zero across participants using a two-sided one-sample t-test. For the ROI-wise cosine-similarity analysis, cosine similarity was tested against zero across participants using two-sided one-sample t-tests, with *P* values adjusted for multiple comparisons using the Benjamini–Hochberg FDR procedure (**Extended Data Fig. 2b**). For the subcategory-preserving shuffle analysis, statistical significance was assessed using a two-sided permutation test. For each permutation, word-pair correspondences were shuffled within semantic relation categories while preserving the source and target subcategories, and the same shuffle was applied across all 128 comparisons (ROIs × relation groups). *P* values were obtained from B = 10,000 permutations and adjusted using the Benjamini–Hochberg FDR procedure.

### Data availability

The fMRI data analysed in this study are publicly available from OpenNeuro under accession code ds003020 (ref. 25). Analyses used data from participants UTS01–UTS09 from version 3.1.1 of this dataset. The processed data generated in this study, including the word representations and relation-vector data used for the analyses, will be made publicly available in a public repository upon publication.

### Code availability

The analysis code used in this study will be made publicly available in a public repository upon publication.

