## Extended Data Figures for "AI-like parallelogram geometry of word relations in the human neocortex"

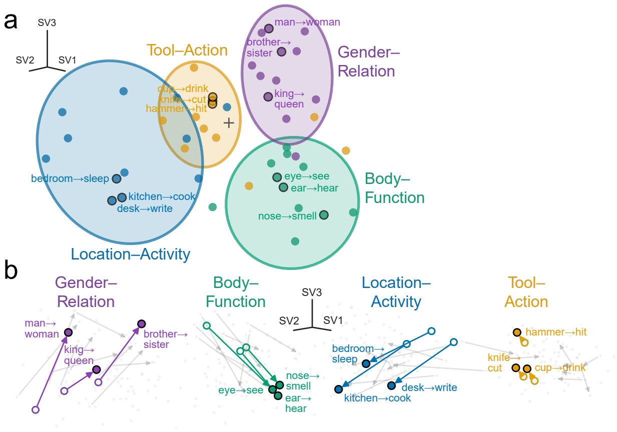


**Extended Data Figure 1. Visualization of relation-vector geometry across repeated cross-validation.** Linear SVMs were trained and evaluated in 100 randomized repetitions of stratified four-fold cross-validation. In each repetition, to balance the sizes of the training and test sets across relation groups, we randomly selected 44 word pairs from the 47 pairs, so that each relation group contributed 3, 3, 3, and 2 pairs to the four folds. **a**, Clusters of recentered relation vectors, projected onto three averaged cross-validated SVM axes (SV1–SV3). More precisely, in each cross-validation fold, the held-out relation vectors were projected onto the three axes of the linear SVM trained on the remaining folds. For each word pair, the resulting projections were averaged across all repetitions in which the pair appeared in the test set. **b**, Parallelogram geometry in word representation space. The three axes are the same as in **a**, but word representations are visualized without recentering relation vectors. The three pairs corresponding to **Fig. 1c,d** are colored.

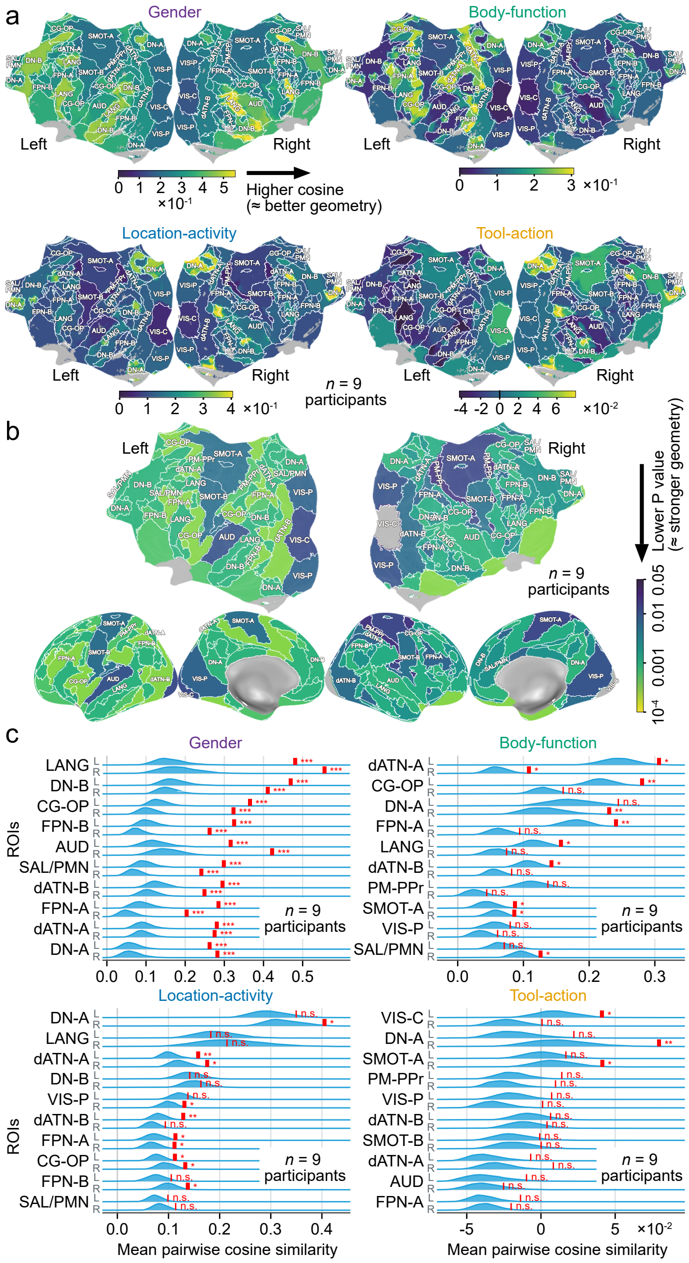


**Extended Data Figure 2. Regional relation-vector parallelism quantified by cosine similarity and tested by subcategory-preserving shuffling.** Parallelism was quantified as the mean pairwise cosine similarity of relation vectors in PCA space and averaged across nine participants. **a**, Relation-vector cosine similarity maps for each relation group. Mean pairwise cosine similarity is shown on flattened cortical maps. **b**, FDR-adjusted P values for gender-relation cosine similarity. Significance is displayed on flattened and partially flattened cortical surfaces (n = 9 participants; two-sided one-sample t-test against zero, df = 8; Benjamini–Hochberg FDR correction across all 128 comparisons [ROIs × relation groups]). Gray, FDR-adjusted P ≥ 0.05. **c**, Word-pair shuffle test while preserving subcategories. For each ROI and relation group, the distribution of cosine similarity for shuffled vectors obtained by subcategory-preserving shuffling (blue) and the cosine similarity for the original vectors (red line) are shown (n = 9 participants; two-sided permutation test, B = 10,000; identical within-subcategory pair shuffles across all 128 comparisons [ROIs × relation groups], with Benjamini–Hochberg FDR correction). For each relation group, ROIs were ordered according to the original cosine similarity in the left hemisphere. ***, FDR-adjusted P < 0.001; **, 0.001 ≤ FDR-adjusted P < 0.01; *, 0.01 ≤ FDR-adjusted P < 0.05; n.s., FDR-adjusted P ≥ 0.05.
